# DNA damage primes hematopoietic stem cells for direct megakaryopoiesis

**DOI:** 10.1101/2023.05.13.540665

**Authors:** Corey M. Garyn, Oriol Bover, John W. Murray, Ma Jing, Karen Salas-Briceno, Susan R Ross, Hans-Willem Snoeck

## Abstract

Hematopoietic stem cells (HSCs) reside in the bone marrow (BM), can self-renew, and generate all cells of the hematopoietic system.^1^ Most hematopoietic lineages arise through successive, increasingly lineage-committed progenitors. In contrast, megakaryocytes (MKs), hyperploid cells that generate platelets essential to hemostasis, can derive rapidly and directly from HSCs.^2^ The underlying mechanism is unknown however. Here we show that DNA damage and subsequent arrest in the G2 phase of the cell cycle rapidly induce MK commitment specifically in HSCs, but not in progenitors, through an initially predominantly post-transcriptional mechanism. Cycling HSCs show extensive replication-induced DNA damage associated with uracil misincorporation *in vivo* and *in vitro*. Consistent with this notion, thymidine attenuated DNA damage, rescued HSC maintenance and reduced the generation of CD41^+^ MK-committed HSCs *in vitro*. Similarly, overexpression of the dUTP-scavenging enzyme, dUTPase, enhanced *in vitro* maintenance of HSCs. We conclude that a DNA damage response drives direct megakaryopoiesis and that replication stress-induced direct megakaryopoiesis, at least in part caused by uracil misincorporation, is a barrier to HSC maintenance *in vitro*. DNA damage-induced direct megakaryopoiesis may allow rapid generation of a lineage essential to immediate organismal survival, while simultaneously removing damaged HSCs and potentially avoiding malignant transformation of self-renewing stem cells.

---

Although HSCs are functionally heterogeneous in terms of lineage output, platelets are the only lineage invariably reconstituted by all HSCs in single-cell transplantation assays.^3, 4^ Furthermore, whereas most hematopoietic lineages arise through successive, lineage-committed progenitors, MKs can derive rapidly and directly from HSCs,^5, 6^ a process stimulated by type I interferon (IFN-I) signaling.^7^ As platelet bias increases with age^8^ and as DNA damage associated with replication stress is involved in HSC aging,^9^ we hypothesized that DNA damage might be a driver of direct megakaryopoiesis.

We first examined whether DNA damage and MK commitment in HSCs are associated *in vivo*. Within 24hrs, irradiation increased HSC size as measured by forward scatter (FSC), induced expression of MK markers (CD41, CD9 and CD62P) **(Extended Data 1a,b),** and increased the fraction of HSCs expressing GATA1, an essential MK transcription factor **(Extended Data 1c,d)**.^2^ Furthermore, the GATA1 expression in individual GATA1^+^ HSCs increased, indicating that these findings were not merely due to selective survival of already MK-committed HSCs **(Extended Data 1d)**. Type I Interferon (IFN-I) signaling, which promotes direct megakaryopoiesis, also recruits HSCs into cell cycle.^10^ After administration of poly-IC, which induces an IFN-I response, deposition of the DNA damage marker, phosphorylated histone H2A.X (γH2A.X),^11^ and the fraction of HSCs positive for the replication stress marker, RPA32,^12^ were increased **(Extended Data 1e,f)**. Together, these data indicate an association between DNA damage and MK commitment *in vivo*. We note that in even in control cells, we observed some staining for γH2A.X **(Extended Data 1e)**, which coincided with nucleoli (not shown). Nucleolar γH2A.X had been observed before in aged HSCs as well,^9^ and its significance is unclear.

To examine the effect of DNA damage on MK commitment *in vitro*, we cultured HSCs (sort gates **Extended Data 2**) for 48hrs in the presence and absence of the topoisomerase 2 inhibitor, idarubicin, which induces double strand breaks.^13^ Idarubicin induced MK markers, increased cell size (**Fig. 1a,b**), and increased the fraction of GATA1^+^ HSCs and GATA1 expression in GATA1^+^ HSC (**Fig. 1c-e**). Idarubicin upregulated some mRNAs encoding MK markers, but not *Gata1* mRNA (**Fig. 1f**), suggesting predominantly post-transcriptional regulation, similar to IFN-I-induced direct megakaryopoiesis.^7^ Hyperploidy in MKs arises either through endomitosis, where cell cycle progresses to the mitotic furrow followed by abortion of mitosis,^2, 14^ or through endocycling, where cells arrest in the G2 phase of the cell cycle and reinitiate DNA synthesis.^15^ As idarubicin induced G2 arrest (**Fig. 1g,h**), and as the G2 phase is a critical checkpoint for polyploidization, we examined the role of G2 arrest. CDK1 is essential for G2 transition, mediated through phosphorylation and subsequent degradation of the phosphatase, CDC25C, by the kinases ATM and ATR. Inhibition of CDC25C prevents removal of inhibitory CDK1 phosphorylation deposited by the kinase, WEE1.^16^ A WEE1 inhibitor, which constitutively activates CDK1,^17^ reduced proliferation and the fraction of cells in G2 (**Fig. 1i-k**), and attenuated the increase in CD9, CD62p and cell size induced by idarubicin, indicating a critical role for DNA-damage induced G2 arrest (**Fig. 1l**). CD41 was not affected however, suggesting that DNA damage alone is sufficient to induce CD41 (**Fig. 1l**).

**Figure 1:**
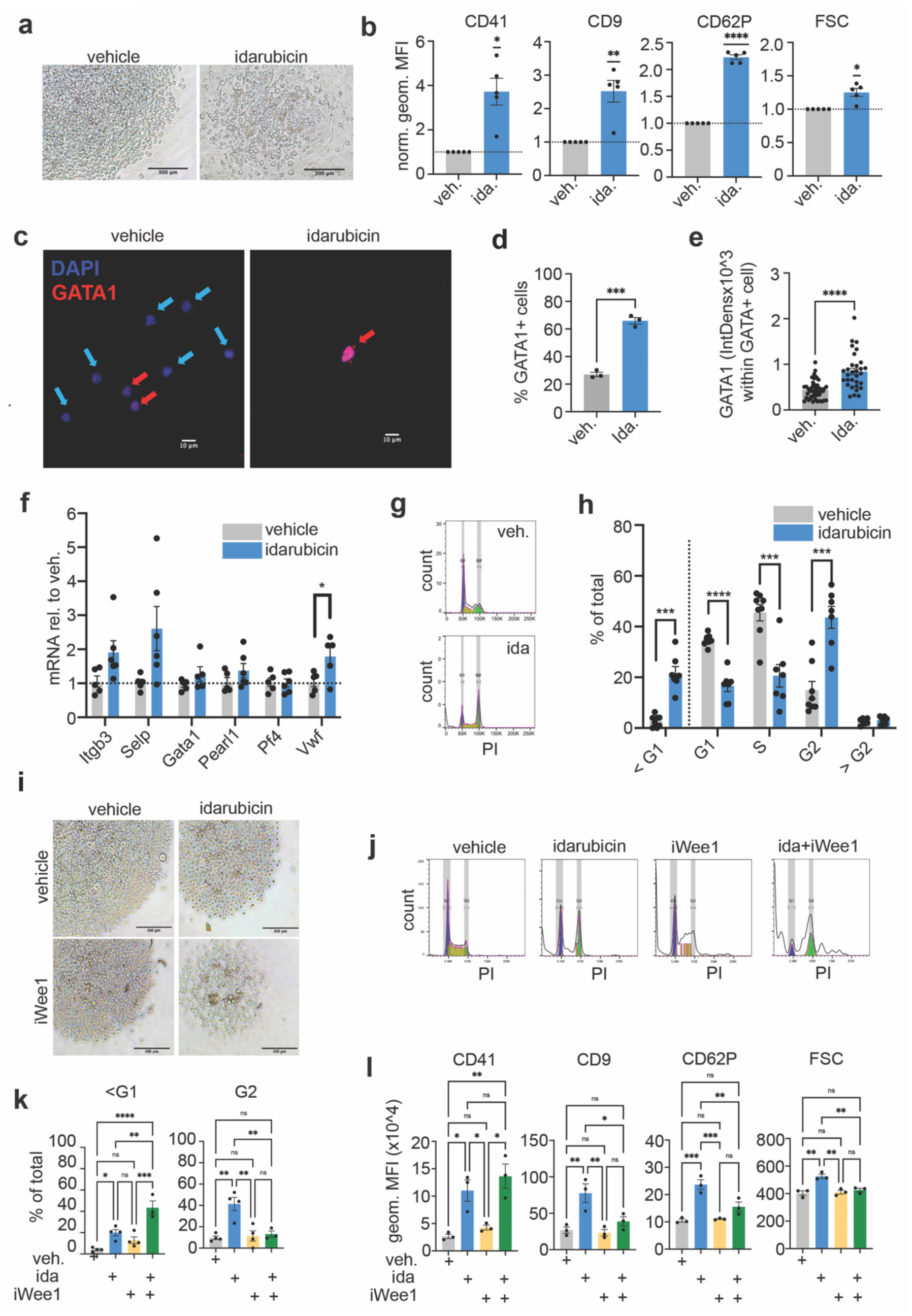
DNA damage induces G2 arrest and a MK phenotype in HSCs. **a**. Brightfield images of vehicle– and idarubicin-treated 48hr HSC cultures (representative of 5 experiments). **b**. Flow cytometric analysis of MK surface markers in 10nM idarubicin– or DMSO vehicle-treated 48hr-HSC cultures (n=5, each averaged across 3 technical replicates and normalized to vehicle, one-sample t-test). **c**. Confocal images of HSCs stained for GATA1 and DAPI after 48hr-culture with vehicle– or idarubicin-culture (representative of 3 experiments). **d**. Quantification of % GATA1 cells after 48hr-culture of HSCs with vehicle– or idarubicin (n = 3, mean +/– SEM, unpaired t-test). **e**. Quantification of GATA1 IntDens within GATA1^+^ population (n = 3, mean +/– SEM, unpaired t-test). **f**. RT-qPCR for MK markers in 48hr vehicle– and idarubicin–treated HSC cultures (normalized to vehicle, n=5, unpaired t-test). **g**. Flow cytometry histograms of cell cycle/ploidy in 48hr HSC cultures with vehicle or idarubicin (representative of 5 experiments). **h**. Quantification of cell cycle/ploidy analyses in 48hr HSC cultures with vehicle or idarubicin (n= 5, mean +/– SEM, paired t-test). **i**. Representative brightfield images of 48hr HSC cultures treated with 10nM Idarubicin and/or 500nM adavosertib (iWee1) (representative of 3 experiments). **j**. Flow cytometry histograms of PI cell cycle and ploidy analyses in 48hr HSC cultures with vehicle, idarubicin and/or adavosertib (representative of 3 experiments). **k**. Quantification of fraction sub-G1 and G2 cells in 48hr HSC cultures with vehicle, idarubicin and/or adavosertib (n=3, one-way ANOVA with Tukey’s multiple comparisons test). **l**. Flow cytometric analysis of MK surface markers in 48hr HSC cultures with vehicle, idarubicin and/or adavosertib (n=3, each experiment averaged across technical replicates, one-way ANOVA with Holm-Sidak’s multiple comparisons). *p ≤ 0.05, **p ≤ 0.01, ***p ≤ 0.001, and ****p ≤ 0.0001.

We next examined the effect of CDK1 blockade. Culture in the presence of the CDK1 inhibitor, RO-3306,^18^ for 48hrs increased cell size, MK marker expression **(Extended Data 3a,b)**, the fraction of GATA1^+^ HSCs and GATA1 expression in GATA1^+^ HSCs **(Extended Data 3c-e)**, and induced G2 arrest and hyperploidy **(Extended Data 3f,g)**. *Gata1* mRNA, however, was not affected, again indicating post-transcriptional regulation **(Extended Data 3h)**. At 96hrs, expression of MK markers (**Fig. 2a**, **Extended Data 3i)**, changes in cell size (**Fig. 2b,c**, **Extended Data 3i)** and induction of MK lineage-associated mRNAs, at this time point including *Gata1* mRNA (**Fig. 2d**), became more pronounced. Hyperploid cells then reached ∼40% and diploid cells disappeared (**Fig. 2e,f**). In multipotential progenitors (MPPs), however, RO-3306 induced apoptosis, attesting to distinct cell cycle regulation in both populations **(Extended Data 3j-l)** and indicating that G2 arrest induces MK specification specifically in HSCs. As these results could be due to the presence of thrombopoietin (TPO), a cytokine that drives both HSC maintenance^19^ and MK development,^20^ we examined the role of TPO. RO-3066 induced MK markers and hyperploidy to the same extent with kit ligand (KL) alone as with KL and TPO (**Fig. 2g,h**). TPO is therefore not required for commitment to the MK lineage induced by a DNA damage response (DDR), potentially explaining why *Tpo^-/-^* mice still have platelets, albeit reduced compared to wt mice.^2^ To demonstrate that HSCs exposed to RO-3066 are truly MK-primed, we used Kusabiro Orange (KuO) mice, where all hematopoietic lineages, including erythrocytes and platelets, express KuO and can be detected in recipients.^21^ While culture in RO-3066 attenuated overall repopulation capacity 12 days after transplantation, the contribution to the platelet lineage was threefold higher compared to vehicle-treated cells, demonstrating MK lineage bias and the expense of overall HSC function (**Fig. 2i,j**).

**Figure 2:**
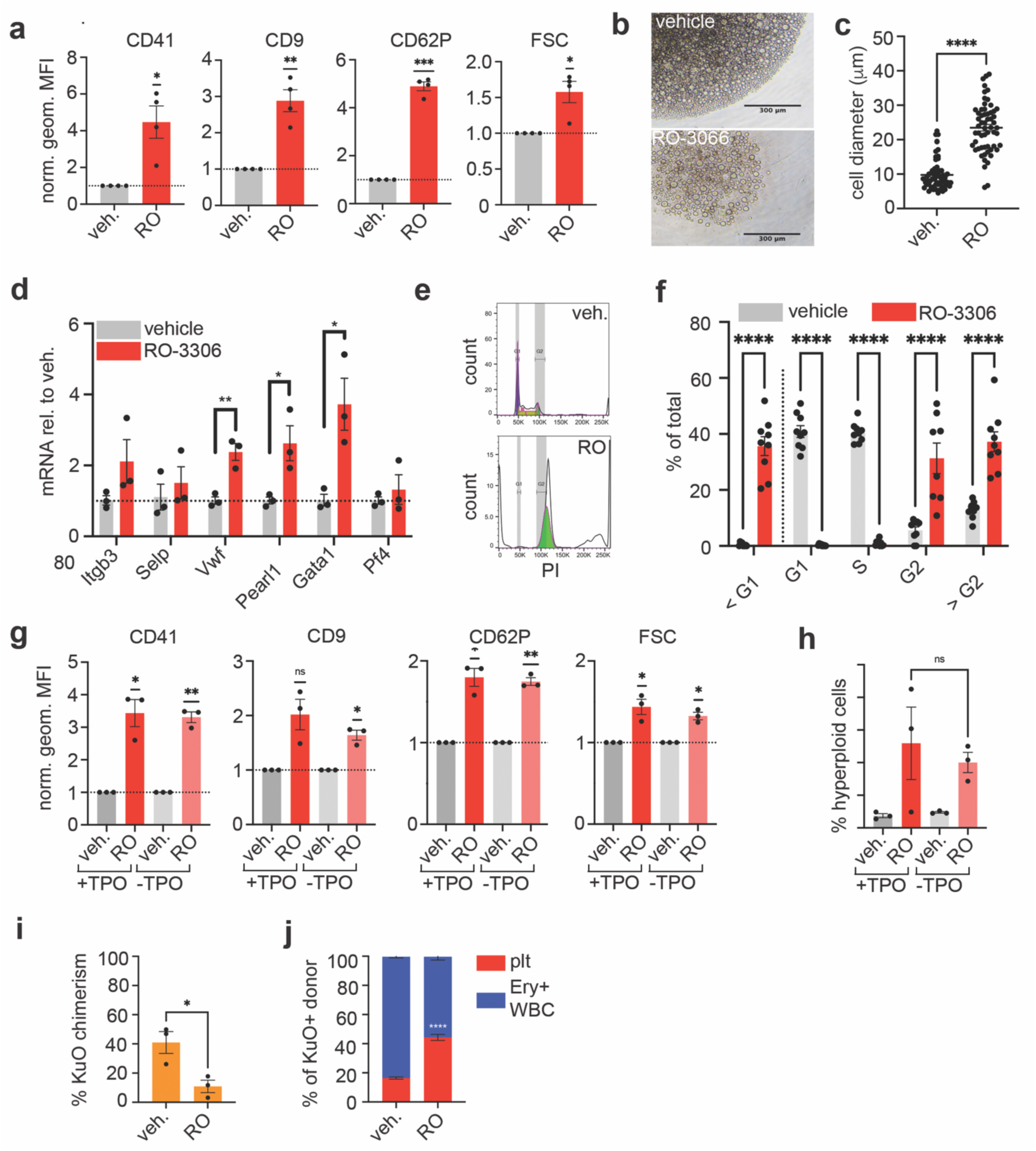
G2 arrest is sufficient to induce direct megakaryopoiesis from HSCs. **a**. Expression of MK surface markers CD41, CD9, and CD62p in 20μM RO-3306– or DMSO vehicle-treated 96hr HSC cultures (n=4, each experiment averaged across technical replicates and normalized to vehicle, one-sample t-test). **b**. Brightfield images of DMSO vehicle– and RO– 3306-treated 96hr HSC cultures (representative of 4 experiments). **c**. Quantification of cell diameter from brightfield images of 48hr HSC cultures treated with vehicle or 20μM RO-3306 (n=1, Mann-Whitney test). **d**. RT-qPCR for MK markers in 96hr vehicle and RO-3306-treated HSC cultures (n=3, normalized to vehicle only, unpaired t-test). **e**. Flow cytometry histograms of cell cycle/ploidy in 96hr vehicle or RO-3306-treated HSC cultures (representative of 4 experiments). **f**. Quantification of cell cycle/ploidy in 96hr vehicle and RO-3306-treated HSC cultures (n=4, >1 technical replicate per experiment, multiple Mann-Whitney tests). **g**. Flow cytometric analysis of MK surface markers in 20µM RO-3306-or vehicle-treated 48hr HSC cultures in with or without TPO (n=3, each experiment averaged across technical replicates and normalized to vehicle, one– sample t-test). **h**. Quantification of % > G2 polyploid cells from vehicle and RO-3306-treated HSC cultures with or without TPO (n = 3, mean +/– SEM, unpaired t-test comparing +/-TPO +RO-3306 conditions). **i**. Quantification of % KuO^+^ PB chimerism 12-days post-transplantation (n = 3 1 recipient mouse per condition and per experiment, unpaired t-test). **j**. Quantification of KuO^+^ platelet vs RBC+PBMC ratio in recipients of DMSO– and RO-3066-treated HSCs (n = 3 recipient mice per condition, unpaired t-test on % KuO^+^ platelets). *p ≤ 0.05, **p ≤ 0.01, ***p ≤ 0.001, and ****p ≤ 0.0001.

MK-biased, CD41^+^ HSCs increase with age and with increasing self-renewal divisions.^8, 22, 23^ Though MK marker expression was lower in CD41^-^ than in CD41^+^ HSCs (sort gates **Extended Data 4a**), RO-3306 caused a similar relative increase in MK marker expression in CD41^-^ and CD41^+^ HSCs (**Fig. 3a**). G2 inhibition-induced MK differentiation is therefore not limited to CD41^+^, MK-primed HSCs. As expected, GATA1 was expressed in a large fraction of CD41^+^ HSCs, but only by few CD41^-^ HSCs (**Fig. 3b**). However, although γH2A.X deposition was on average similar in the CD41^+^ and CD41^-^ populations (**Fig. 3c**), the rare GATA1^+^ cells within the CD41^-^ HSC population showed strikingly elevated γH2A.X deposition (**Fig. 3d,e**, **Extended Data 4b)**. These data suggest that a DDR is associated with GATA1 expression in CD41^-^ HSCs prior to transition to CD41^+^ HSCs and suggest DNA repair in the latter. This notion was supported by the finding that RO-3066 reduced γH2A.X (**Fig. 3f,g**).

**Figure 3:**
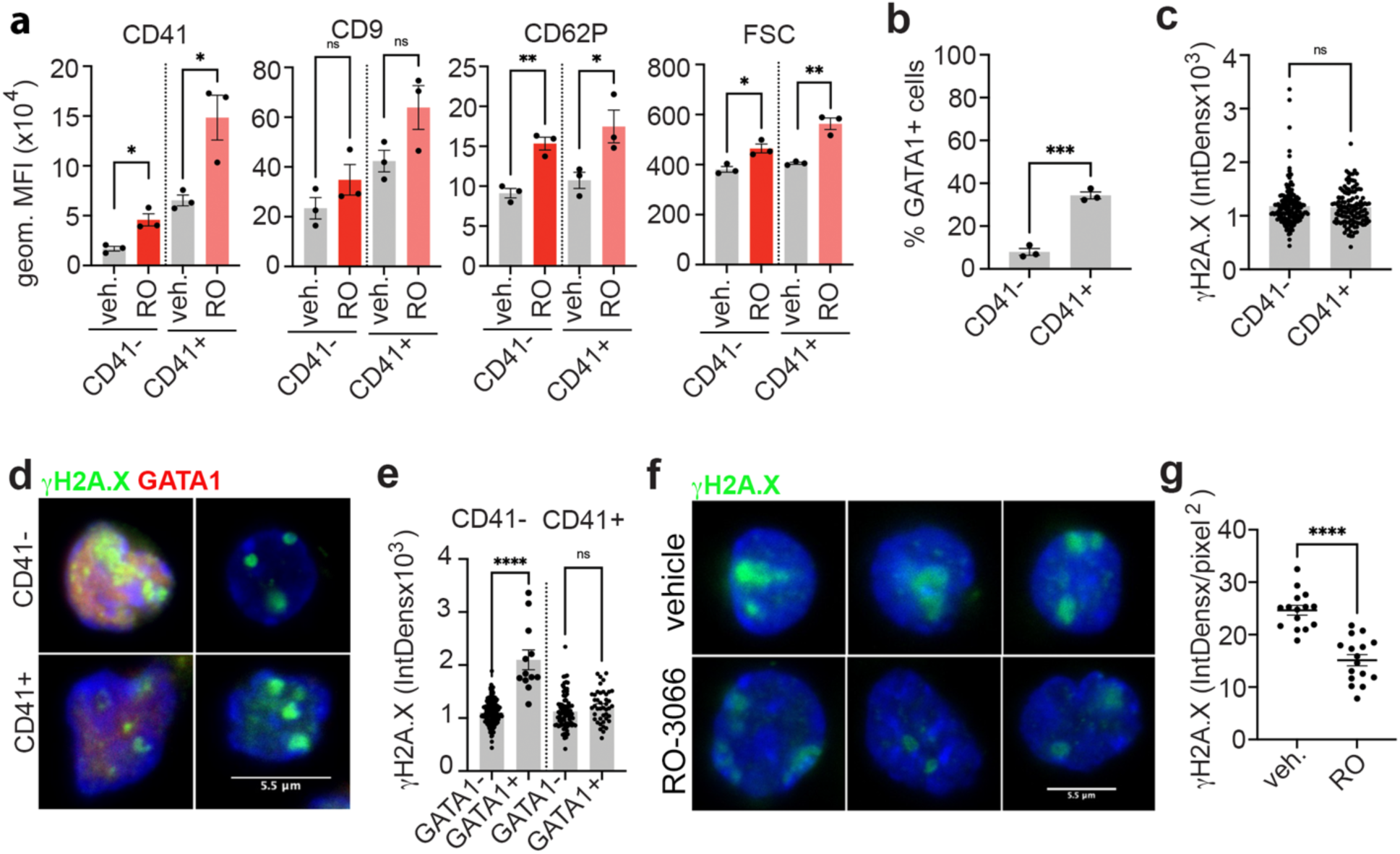
DNA damage induces GATA1 protein expression and differentiation to a CD41^+^HSC pool. **a**. Expression of MK surface markers in RO-3306– or DMSO vehicle-treated 48hr cultures of CD41^+^ and CD41^-^ HSCs (n=3, each experiment averaged across technical replicates, unpaired t–test). **b**. Fraction GATA1^+^ cells in CD41^-^ and CD41^+^ HSCs (n=3, unpaired t-test). **c**. γH2A.X integrated density in fresh CD41^-^ and CD41^+^ HSCs (n=3, mean +/– SEM, Mann-Whitney test). **d**. Confocal images of fresh CD41^-^ and CD41^+^ HSCs stained for γH2A.X and GATA1 (representative of 3 experiments). **e**. Quantification of γH2A.X in GATA1^+^ and GATA1^-^ CD41^-^ and CD41^+^ HSCs (n=3, unpaired t-test for CD41^-^ HSCs, and Mann-Whitney tests for CD41^+^ HSCs). **f**. Confocal images of 48hr HSC cultures in 20µM RO-3306 and DMSO control, stained for γH2A.X (green) (n=1). **g**. Quantification of nuclear γH2A.X IntDens normalized to cell area (pixel^2^) based on DAPI stain (n=1, Mann-Whitney test). *p ≤ 0.05, **p ≤ 0.01, ***p ≤ 0.001, and ****p ≤ 0.0001.

As a DDR induces MK commitment, and as HSCs are prone to replication stress, we hypothesized that MK commitment may be a barrier to HSC maintenance *in vitro*, which remains challenging. 7d-culture in previously optimized conditions for HSC maintenance^24^ caused a striking increase in γH2A.X deposition **(Extended Data 5a,b)**, comet assay tail moment **(Extended Data 5c)** and 53BP1 staining **(Extended Data 5d),** indicative of increased DNA damage. Abundant deposition of RPA32 indicated damage caused by replication stress **(Extended Data 5f)**.

One cause of replication stress is misincorporation of uracil into DNA,^25^ the removal and repair of which provokes a DDR.^26^ Incorporated uracil is rapidly removed primarily by uracil N-DNA glycosylase (UNG1/2).^27^ We observed UNG2 to be strikingly present in nucleoplasm and nucleoli of HSCs and MPPs, and to more variable extent in CMPs, but absent from lin^+^ cells (sort gates **Extended Data 2),** indicating that HSCs and MPPs are poised to excise uracil (**Fig. 4a**, **Extended Data 6a,** *Ung*^-/-^ control **Extended Data 6b)**. Semi-quantitative measurement of uracil content^28^ in fresh and cultured *Ung^-/-^*cells (classically used because of rapid removal of uracil in wt cells)^29^ showed strikingly increased uracil incorporation in cultured HSCs, but not cultured BM-derived macrophages (**Fig. 4b**). Importantly, administration of polyI:C *in vivo*, which results in replication stress **(Extended Data 1e,f)**, induced striking uracil misincorporation in HSCs, but not in lin^+^ cells (**Fig. 4c**). Cycling HSCs are therefore prone to uracil misincorporation, *in vivo* and *in vitro*.

**Figure 4:**
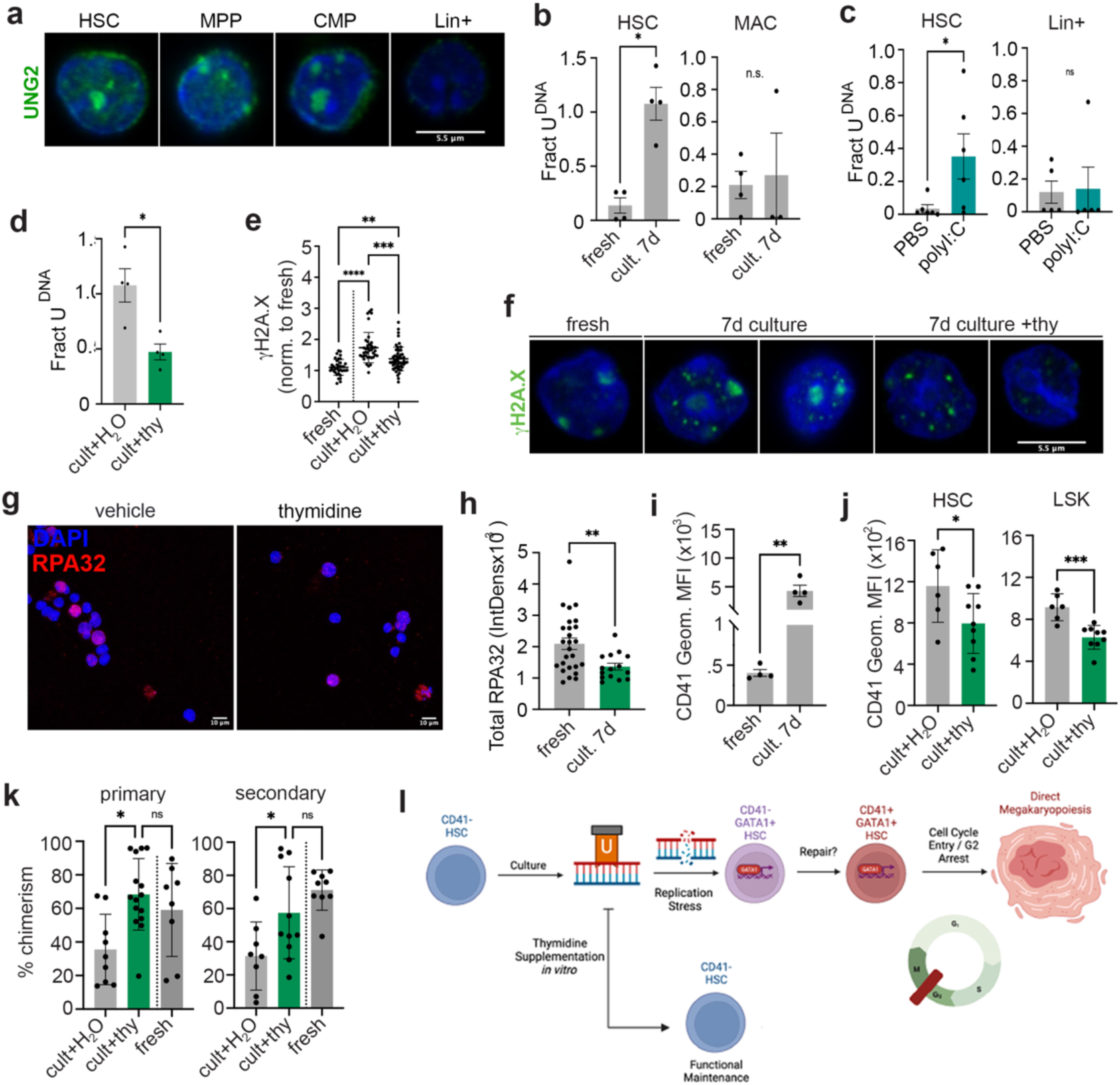
Uracil misincorporation induces DNA replication stress and impairs HSC functional maintenance *in vitro*. **a**. Confocal images of BM populations stained for UNG2 and DAPI (n=1). **b**. Quantification of uracil DNA content (Frac U^DNA^) in fresh and 7-day cultured *Ung^-/-^* HSCs and macrophages (MAC) (n = 3, unpaired t-test). **c**. Frac U^DNA^ in HSCs and lin^+^ cells 24hrs after administration of polyI:C *in vivo* (n=6, unpaired t-test). **d**. Frac U^DNA^ in 7-day cultured *Ung-/-* HSCs with or without 100µM Thy. (n = 3, unpaired t-test). **e**. Quantification of nuclear γH2A.X in fresh and 7-day resorted HSCs from culture with and without thymidine (n = 3, one-way ANOVA). **f**. Confocal images of fresh or 7-day resorted HSCs stained for γH2A.X and DAPI (representative of 3 experiments). **g**. Confocal images of 7-day HSC cultures with or without thymidine stained for RPA32 and DAPI (n = 1). **h**. Quantification of RPA32 IntDens in 7-day with or without thymidine (n = 1, unpaired t-test). **i**. CD41 geometric MFI in fresh and 7-day cultured phenotypic HSCs (n = 3, unpaired t-test). **j**. CD41 geometric MFI of HSCs and LSKs after 7-day culture with or without thymidine (n = 3, unpaired t–tests). **k**. Quantification of 16-week primary and secondary post-transplantation chimerism of fresh HSCs and HSCs cultured with or without thymidine for 14d (n = 3, one-way ANOVA). **m**. Model of induction of direct megakaryopoiesis by uracil misincorporation and DNA damage. *p ≤ 0.05, **p ≤ 0.01, ***p ≤ 0.001, and ****p ≤ 0.0001.

As thymidine competes with deoxyuridine, we examined its effect on uracil incorporation and HSC function. Thymidine induces G1 arrest.^30^ High-dose (1mM) thymidine was indeed toxic (not shown). A lower concentration (100μM), however, did not induce G1 arrest in HSCs, but caused complete arrest in MPPs **(Extended Data 6c,d)**, again attesting to differential cell cycle regulation in HSCs and MPPs. Thymidine reduced uracil incorporation (**Fig. 4d**), γH2A.X (**Fig. 4e,f**) and RPA32 (**Fig. 4g,h**) deposition in HSCs cultured for 7 days, and concomitantly attenuated induction of CD41 in HSCs and in progenitors (**Fig. 4i,j**, **Extended Data 6e)**. Thymidine furthermore increased competitive repopulation capacity in primary and secondary transplantations after 14d culture, such that net maintenance was achieved (**Fig. 4k**). Limiting dilution analysis indicated 20–fold better maintenance of functional HSCs after 14d culture in the presence of thymidine compared to vehicle **(Extended Data 6f,g)**. Using donor HSCs from KuO mice, addition of thymidine to 14-day cultures increased long-term repopulation in all lineages, including the erythroid and MK lineages **(Extended Data 6h)**, indicating maintenance of multipotential HSCs. Next, we overexpressed dUTPase (DUT, encoded by *Dut*), which generates dUMP from dUTP, thus providing a substrate for dTMP and dTTP synthesis. Lentiviral transduction of DUT followed by 7-day culture increased long-term repopulation capacity **(Extended Data 6i)**, providing further evidence for a role of uracil incorporation in the decline in HSC function *in vitro*. Together, these data show that thymidine reduces uracil misincorporation and in doing so attenuates DDR-induced MK commitment and rescues maintenance of multipotential HSCs.

We conclude that a DDR in HSCs promotes direct megakaryopoiesis at least in part through post-transcriptional mechanisms driven by G2 arrest (**Fig. 4l**). In addition to endowing HSCs with the capacity to rapidly generate short-lived elements essential for immediate organismal survival, this mechanism may protect the integrity of the HSC compartment, as DNA-damaged HSCs are shunted to MK lineage, thus removing them from the HSC compartment. Furthermore, the accumulation of MK-primed HSCs with age^22^ may be caused by age-related replication stress.^9^ Importantly, our data indicate that DNA damage through replication stress is caused, among others factors, by uracil misincorporation, and that ensuing direct megakaryopoiesis is a barrier to *in vitro* maintenance of HSC function.

## Acknowledgments

This work was supported by grants R01HL135039 (HWS) and NIH S10 OD032447 (HWS). The authors thank Dr. Shan Zha (Herbert Irving Comprehensive Cancer Center, Columbia University Irving Medical Center) for advice.

## Author Contributions

CMG performed most experiments and contributed to concept, OBC performed staining for nucleolar proteins and comet assay, JM oversaw image quantification and wrote macro for GATA1 quantification, MJ, KJSB and SRS performed uracil determination, HWS provided concept, supervised and wrote the manuscript.

## Methods

### Mice

C57BL/6 (CD45.2) and PepboyJ (CD45.1) mice were either purchased from Jackson Laboratories (#000664; 002014) or bred in-house. Kusabira Orange (KuO+) mice were gifted by Dr. Nakauchi (University of Tokyo) and rederived from embryos in maximum barrier conditions through Charles River Laboratories (location). All mice were housed in Helicobacter– and Pasteurella-free rooms, and fed and watered *ad libitum*. All mouse holding and experimental techniques were performed in accordance with approved protocol AAABM6567 from the Columbia University Institutional Animal Care and Use Committee (IACUC).

## Mouse hematopoietic tissue preparation for cell sorting

BM cells were isolated from hindlimbs by crushing in 1× PBS in a non-tissue culture treated petri dish. Cell suspensions were centrifuged at 420 x g for 3 minutes at room temperature, then resuspended in 1× ACK lysis (Gibco) and incubated for 5 minutes RT to lyse red blood cells. ACK lysis was neutralized with 1× PBS and samples were again centrifuged before proceeding to FACS.

## Mouse hematopoietic cell purification by fluorescence-activated cell sorting (FACS)

Hematopoietic stem and progenitors were enriched prior to sorting by staining with anti cKit-biotin antibody (clone 2B8; 1:100 dilution) and positive selection using magnetic anti-biotin microbeads on LS columns (Miltenyi Biotec., Gaithersburg, MD). For HSC isolation, cKit-enriched BM and unenriched control BM were stained for Lineage markers (FITC-conjugated CD2, CD3, CD4, CD8a, CD11b, CD19, B220, Ter119, Ly6-C/Ly6-G (Gr1), all 1:200), CD48-A700 1:200, Sca1-PB 1:200, cKit-APC-Cy7 1:200 (clone ACK2), CD150-PE-Cy7 1:100, FLT3-PE or FLT3-APC 1:200, and, where noted, CD41-PE 1:400 (see **table S1** for list of antibodies). Cells were stained with surface markers for 30 minutes at room temperature protected from light. Populations were sorted on a FACSAriaII (BD Biosciences, Franklin Lakes, NJ) by selecting for the following surface marker phenotypes: HSCs were Lineage^lo/-^cKit^+^Sca1^+^CD48^-^Flt3^-^CD150^+^ and CD41^+/-^ when noted; MPPs were Lineage^lo/-^cKit^+^Sca1^+^CD48^+^Flt3^+^; CMPs were Lineage^lo/-^cKit^+^Sca1^-^; and mature blood cells were sorted as Lineage+.

## Culture of mouse hematopoietic stem cells

Purified HSCs were centrifuged into 1.5ml Eppendorf tubes coated with MACS buffer and containing complete mouse HSC culture (described below). Sorted HSCs were then resuspended in fresh complete mouse HSC media without calcium before plating at 2000-3000 cells per well in 96-well round bottom plates without TC treatment. Mouse Basal culture media was made from Ca2+-free DMEM (US Biologicals) reconstituted with 4.5g/L glucose and 1.1g/L sodium bicarbonate at pH 7.4. Mouse complete HSC culture media was prepared with Ca2+-free basal media supplemented with 200ng/ml rmSCF (Peprotech, Cranbury, NJ), 30ng/ml rmTPO (Peprotech), 1× StemPro34 supplement (Life Technologies, Carlsbad, CA), 1× Glutamax (Thermo Fisher Scientific, Waltham, MA), 1× Pen/Strep (Life Technologies), and 10mM HEPES buffer. Complete HSC culture media was prepared with either 2.00mM or 0.02mM CaCl_2_. HSCs were cultured in various media conditions in 37°C 5% O_2_ 5% CO_2_ incubators. Half-media changes were performed every 48 hours without disturbing the cell pellet, and cultures were passaged 1:2 at 7 days after plating.

Small molecules were added following concentrations (with DMSO vehicle at ≤ 0.1% v/v): 10nM Idarubicin in DMSO (Selleck, Houston, TX), 20µM RO-3306 in DMSO (Selleck), 500nM Adavosertib in DMSO (iWee1; Selleck). At each analysis timepoint, brightfield images of cultures are generated on Leica DMI8 Inverted Microscope. Analysis of cell diameter was performed by setting a scale with a millimeter ruler image, and using the measurement tool to calculate the length in μm along the longest diameter of individual cells.

## Flow cytometric analysis of cultured hematopoietic stem cells

At designated timepoints, cells were transferred to 5ml polystyrene flow tubes, washed with 1ml 1× PBS and centrifuged at 420 x g for 3 minutes. Cells were resuspended in 100µl MACS buffer consisting of 1% BSA and 10mM EDTA in PBS with the appropriate conjugated surface marker antibody panels depending on the experiment (see **Table S1**). Surface marker staining was performed for 30 minutes at room temperature protected from light, followed by washing with 1ml PBS. Samples were resuspended in 100µl MACS buffer and analyzed using the Novocyte Quanteon Flow Cytometer (Agilent), the BD LSRII (BD), or the BD Fortessa (BD).

## Immunofluorescent staining and imaging

96-well flat-bottom glass plates (Cellvis, Mountain View, CA) were coated with 5µl Cell-Tak (Corning, Corning, NY) and precipitated using 1M NaHCO_3_ for several hours at room temperature. Cell-Tak solution was aspirated, and 1000-5000 fresh or cultured HSCs were plated with gentle centrifugation at 20 x g for 2 minutes. Immunofluorescent staining was performed as follows: 1) fixation at room temperature for 15 minutes in 250µl 4% Paraformaldehyde in PHEM buffer, followed by 2) three washes with PBS, 3) permeabilization with 250µl 0.1% Triton X-100 in PBS for 15 minutes, 4) 3× washes in PBS, and 5) blocking for 1 hour in 100µl of blocking solution containing 1× PBS with 2% BSA 0.1% Triton X-100 and 1:100 CD16/CD32 Mouse Fc block (BD). Pre-extraction for DNA damage marker imaging was performed by permeabilization of live cells in 0.1% Triton X-100 in PHEM buffer for 10 minutes to extrude solubilized proteins while preventing microtubule catastrophe, followed by the same fixation, permeabilization, and blocking protocol described in the normal approach. Primary antibody incubations were performed overnight at 4°C in humidifying chamber with 50µl of the following antibody dilutions in blocking solution: 1:100 mouse-anti-phospho-H2A.X (Ser139, Millipore), 1:50 rat-anti-RPA32/RPA2 (4E4, abcam), 1:50 rat-anti-GATA1 (N6, Santa Cruz, Santa Cruz, CA), 1:500 rabbit-anti-53BP1 (NB100–904, Novus, Englewood, CO), and 1:200 rabbit-anti-UNG2 (pAb, Thermo Fisher). Cells were washed 3× in PBS and incubated for 2 hours at room temperature protected from light in 50ul of 1:500 AlexaFluor secondary antibodies (Thermo) in blocking solution (see **Table S1)**. After three additional washes in PBS, stained cells were mounted in fluorescent mounting media with DAPI (OriGene, Rockville, MD) and stored at 4°C in tinfoil until ready to image.

Cells were imaged at 63× with immersion oil and electronic zoom on a Leica Stellaris 8 confocal or Zeiss LSM 710 confocal microscope. For γH2A.X nucleolar analyses, individual slices in the center of cells were manually selected. For GATA1 nuclear stains, an ImageJ macro was developed, encoding a similar watershedding approach to determine nuclear boundary of z-projected images based on DAPI counterstain, then measuring GATA1 integrated density within the automated boundary and determining GATA1 positivity based on a manually-determined threshold.

## Lentiviral transduction

Lentiviral particles were produced by seeding LentiX-293 cells (Takara, San Jose, CA) at 7×10^5^/cm^2^ in Ultra Culture serum-free media (Lonza, Basel, Switzerland) overnight followed by transfection of each packaging and expression construct (1:1:1) using X-tremeGENE HP DNA transfection reagent (Sigma-Aldrich, St. Louis, MO) for 2 hours. Media from transduced cells were pooled after 36–48h and clarified by centrifugation at 3000 x g for 15min. Resulting viral supernatants were then concentrated by ultracentrifugation (100,000 x g), resuspended in StemPro-34 media and stored at −80°C. Virus titer was calculated from transduction of NIH-3T3 fibroblasts serial dilutions of the viral preparation. Sorted HSCs were transduced with 150-200 MOI lentivirus particles in the presence of 6µg/mL polybrene (Sigma) and spun at 900 x g for 20min at 20°C. Supernatant was aspirated and replaced with complete media and cultured overnight. Transduction efficiency of cells was confirmed after 24hr by flow cytometry against the GFP fluorescent reporter.

## Competitive repopulation

10% percent of final culture volume from 7– or 14-day HSC cultures was mixed with 2×10^5^ BM cells from CD45.1/2 competitors and intravenously transplanted into CD45.2 mice lethally irradiated with a two split doses of ∼450cGy X-ray irradiation. Transplantations were performed at day of culture seeding (day 0) and the endpoint reported for the relevant experiment. For competitive transplantation experiments, at least three independent transplantations, each with at least 4 recipients per condition was performed, and donor chimerism from all recipients was pooled for statistical analysis. Sample sizes were estimated from power calculations based on results of the preliminary experiment. For secondary transplantations, 10^6^ BM cells from primary recipients were transplanted into at least three lethally irradiated CD45.2 secondary recipients.

For KuO^+^ 48-hour mouse HSC transplants, phenotypic KuO^+^ HSCs were seeded at day zero to yield 20-40k cells at end of culture duration in either DMSO vehicle– or 20µM RO-3306 conditions. Final culture cellularity was determined using a hemacytometer. Equal cell numbers were prepared for each culture condition, combined with 5×10^5^ CD45.1/2 whole BM competitor cells, and transplanted into a single CD45.2 mice lethally irradiated as described above.

## Limiting Dilution Assay (LDA)

In LDA experiments, cohorts of CD45.2 recipients received 100, 500, or 1000 cells from donor CD45.1 HSCs after 2 week of culture in 0.02mM Ca2+ complete HSC media +/-100µM thymidine together with 2×10^5^ competitor CD45.1+CD45.2+ BM cells, allowing calculation of HSC frequency based on the number of non-repopulated mice using Poisson statistics (http://bioinf.wehi.edu.au/software/elda/) 16 weeks after transplantation. Recipients showing ≥1.0% CD45.1 donor contribution and contributing to B, T and myeloid lineages were considered positive.

## Peripheral blood preparation and flow cytometric analysis

Peripheral blood was collected by submandibular vein puncture with a 5.5mm sterile disposable lancet into 0.5mM EDTA chelation to prevent coagulation. Red blood cells were lysed once with 5-minute 1× ACK incubation, neutralized with 1× PBS, and centrifuged at 450g for 5min at RT. The resulting pellet was resuspended in antibody staining mix (see **Table S1**) and incubated for 30 minutes at RT protected from light. Stained samples were washed with 1× PBS, resuspended in MACs buffer, and analyzed on either an LSRII (BD), Fortessa (BD), or Novocyte Quanteon. Transplant recipients were analyzed for B-cell, T-cell, and myeloid lineage chimerism based on CD45 allele phenotypes using the surface marker panels outlined in **Table S1**.

For KuO^+^ chimerism in RBC and platelet lineages, 10µl of unlysed, chelated peripheral blood was stained for 30 minutes at RT protected from light for appropriate surface markers (see **Table S1**), then diluted in 1ml MACs buffer and immediately analyzed on a Novocyte Quanteon. Forward and side scatter were collected on the logarithmic scale to separate platelet populations from RBCs and PBMCs based on cell morphology.

## Cell cycle and ploidy analysis

10^3^ –10^4^ cultured HSCs were transferred to 5ml polystyrene flow cytometry tubes and washed twice in 2ml of 1x PBS without calcium/magnesium and centrifuged 5min at 200 x g. After the second wash cells were resuspended in 200µl PBS and fixed by adding 800µl cold 100% ethanol dropwise with gentle vortexing, then stored at –20°C for ≥ 2 hours. Fixed cells were washed twice with 2ml PBS and centrifuged for 5min at 800 x g. Cells were then stained for 20min in 37°C water bath protected from light using 500µl of the following staining solution: 1µg/ml propidium iodide (Thermo), 0.2mg/ml RNAseA (Sigma, St. Louis, MO), and 0.1% Triton X-100 (Sigma) in PBS. Propidium iodide DNA fluorescence was collected on the PE-TR channel on a linear scale using the Novocyte Quanteon Flow Cytometer (Agilent, Santa Clara, CA), and cell cycle stages were determined using the Watson pragmatic model in Prism 9 with defined G1 and G2 ranges.

## cDNA preparation and reverse-transcription quantitative PCR (RT-qPCR)

### RNA purification and DNase treatment

10-20×10^3^ cultured mouse HSCs were collected at either 48– or 96-hours after seeding, lysed in 1:3 Trizol LS (Life Technologies), and stored at –80°C. RNA was extracted from 400μl thawed Trizol sample with 100μl chloroform, vigorous vortexing for 20 seconds, centrifugation at 12,000g for 15 minutes at 4°C, and transfer of aqueous layer to a new tube. RNA was precipitated with 2.5µl of 5mg/ml linear acrylamide (Thermo) and 200μl of 100% isopropanol, vortexing and 5–minute room temperature incubation, and centrifugation at 12,000g for 10 minutes at 4°C. Pelleted RNA was washed with 1ml cold 70% ethanol in nuclease-free water and centrifuged 7500 x g for 5 min 4°C. RNA pellet was isolated by aspirating ethanol and air-drying for 5 min at room temperature. DNase treatment was performed by resuspending the RNA pellet in 51μl nuclease-free water, 3U DNaseI (Invitrogen, Waltham, MA) and 6µl 10x DNase buffer (Invitrogen), vortexing and incubating 15 min at room temperature. DNase treatment was stopped by adding 6μl of 25mM EDTA (Invitrogen) and heating samples to 65°C for 10 minutes. RNA was purified by adding 34μl nuclease-free water, 10µl 3M NaOAc, and 250μl 100% ethanol and incubating – 80°C for > 1 hour. After centrifugation at 7500 x g for 5 min at 4°C no RNA pellet is visible, yet >95% volume is aspirated, and remaining RNA is washed twice with 1ml cold 70% ethanol before air-drying and resuspending in 11µl nuclease-free water.

### Reverse transcription reaction and cDNA preparation

Primer annealing was performed by adding to 11μl purified RNA 1ul 100µM random hexamer primers (Thermo) and 1μl 10mM dNTP mix (Thermo), and incubating 65°C for 5 min on a thermocycler. The reverse transcription reaction was prepared by adding to annealed RNA 4μl 5x First-strand buffer (Invitrogen), 1ul 0.1M DTT (Invitrogen), 40U RNAseOUT recombinant ribonuclease inhibitor (Invitrogen), and 200U Superscript III Reverse Transcriptase (Invitrogen). Reverse transcription was performed in a thermocycler at 25°C for 5 min, followed by 50°C for 60 min, and inactivated at 70°C for 15 min. The final product was diluted 1:10 in nuclease-free water to prevent reactivation of RT enzymes during PCR, and stored at –20°C.

### Reverse transcription-quantitative PCR

Quantitative PCR was performed on a QuantStudio 5 Real-Time PCR System (Invitrogen) with PowerSYBR Green PCR Master Mix (Applied Biosystems, Waltham, MA) or Taqman Probes (see **Table 2**) at 15µl final volume in Microamp Optical 384-well Reaction Plate with Barcode (Applied Biosystems). Target genes were analyzed using primer sets outlined in **Table 2** The reaction conditions were 50°C hold for 2 min, 95°C for 10 min, then 40 cycles of 95°C for 15 second and 60°C for 1 min. Melt curves were immediately determined following PCR using 95°C for 15 seconds, 1.6°C/second decrease to 60°C, then 0.075°C/second increase to 95°C and a final 15 second hold. Gene expression was normalized to *Hprt* or *Gapdh* housekeeping control to determine ΔΔCt between experimental conditions.

## Detection of uracil DNA content

Semi-quantitative detection of uracil content in DNA was performed in collaboration with Dr. Susan Ross and colleagues at University of Illinois at Chicago. *Ung+/+* or *Ung-/-* HSCs or macrophages (CD11b+Gr-1– BM cells) were sorted and immediately analyzed for uracil, or cultured for 7-days in high-calcium culturing conditions and resorted for analysis as described previously. Excision qPCR was used to determine the uracil-containing fraction of DNA as described in Meshesha et al.,^31^ with some modifications. Samples were split into two equal portions, one control sample and one treated with UDG. For UDG-treated sample, 0.125 units of LTR Escherichia coli uracil-DNA glycosylase (New England BioLabs) was added to Promega qPCR master mix to excise uracils from gDNA. The qPCR thermocycler reaction was modified to include the UDG reaction time and heat cleavage of the resulting abasic sites. Reactions were performed with the following cycling protocol: 37°C for 30 min (UDG reaction), 95°C for 5 min (abasic site cleavage), 40 cycles of denaturation at 95°C for 10 sec and annealing and extension at 60°C for 30 sec. 45S rDNA primers were used to amplify DNA **(see** **Table 2)**. Primers targeting *Gapdh* were used to calculate the fraction of uracil-containing DNA (Frac U) using the threshold cycle (ΔΔCT) method.

## Intraperitoneal injections and *ex vivo* analyses

Intraperitoneal injections were performed with 5mg/kg b.w. poly(I:C) in PBS (Invitrogen). For irradiation analyses, mice were exposed to 450cGy X-ray irradiation. Following designated *in vivo* treatment timeframe, HSCs were isolated and sorted from bone marrow using approaches described above.

## Comet assay

The assay was done following the manufacturer’s protocol. Briefly, 2000 HSCs (either freshly sorted or re-sorted from a 14-day culture) cells were mixed with low melting point agarose at a 1:10 dilution and added on top of the slide with a previously polymerized base of agarose. After polymerization of the cell-agarose mixture the slide was put in the lysis buffer overnight at 4 °C. The lysis buffer was replaced with the pre-chilled alkaline solution for 30 mins a 4°C. Electrophoresis was done in cold alkaline solution for 30 min at 15 V (300 mA). The slide was immersed horizontally in cold diH2O for 2 min 3x, then changed to 70% cold ethanol for 5 min. The slide was dried at room temperature for ∼1 hour and stained using the provided dye. Slides were imaged using an epifluorescence microscope and the images were analyzed using the Comet Score software. For the positive control whole bone marrow cells were incubated for 20 min in 0.1% H_2_O_2_.

## Other reagents

All reagents, excluding antibodies and oligonucleotide primers, are listed in **Table S3**.

## Statistical analysis

For all comparisons, normality of data was determined using Shapiro-Wilk normality test. For parametric analysis, unpaired t-tests or one-way ANOVA were performed as indicated in figure legends, while Mann-Whitney and Kruskal-Wallis tests were performed on non-normal data. For MK surface marker phenotypes, geometric MFI fold-change from control conditions was calculated and analyzed using one-sample t-tests. Data are shown as mean±SEM throughout.

All statistical tests were run using Prism 9 software.

31. Meshesha, M. *et al.* Deficient uracil base excision repair leads to persistent dUMP in HIV proviruses during infection of monocytes and macrophages. *PLOS One* **15**, 1–26 (2020).

## Extended Data

**ED1:**
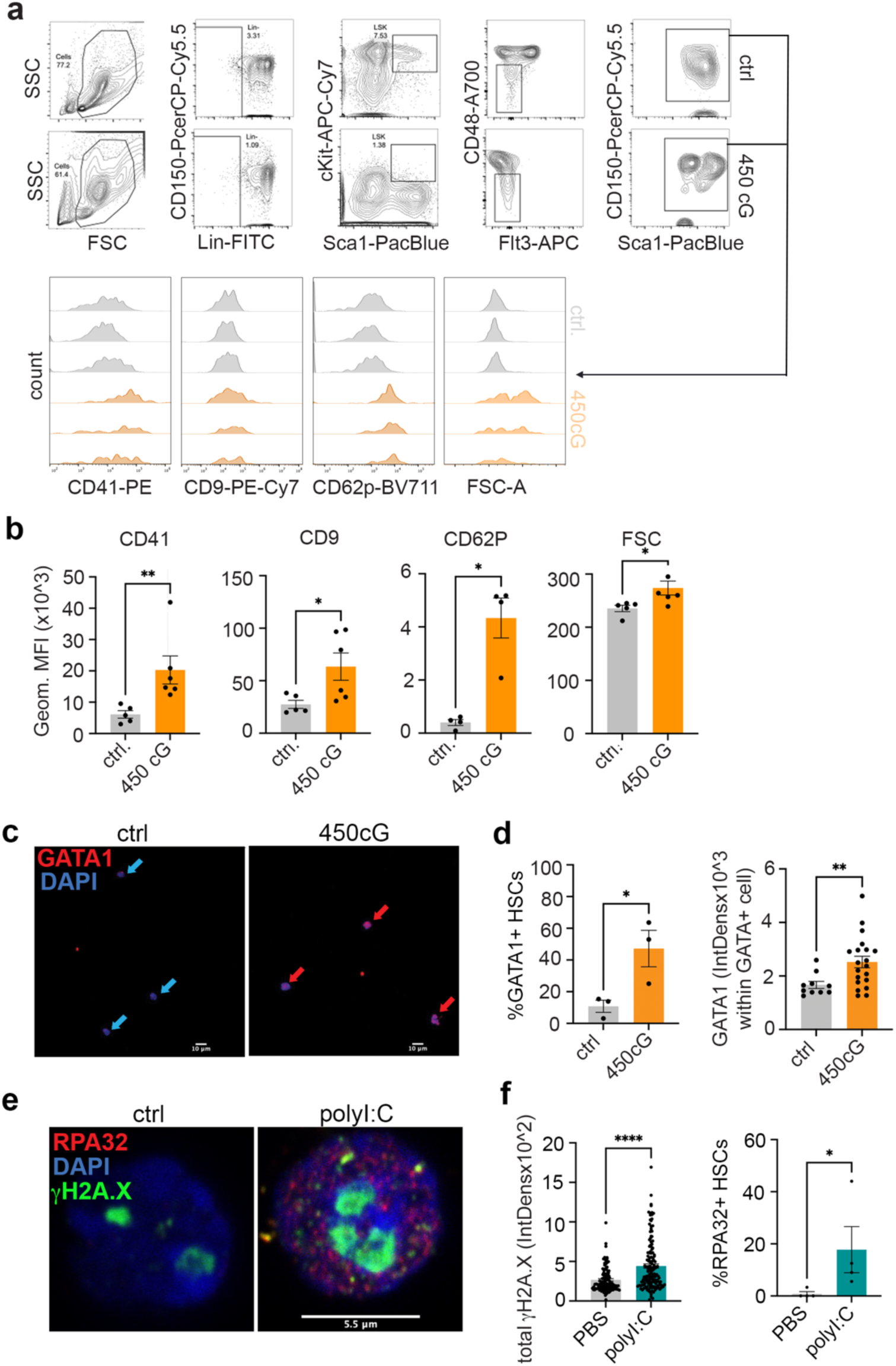
Association of DNA damage, cell cycling and MK marker induction. (a) Representative flow cytometry plots of control and 24hr post-450cGy X-ray-irradiated BM stained for HSC and MK markers (n=3). **(b)** Quantification of **(a)** gated on HSCs (n=3, each experiment averaged across technical replicates, unpaired t-tests (Mann-Whitney for CD9)). **(c)** Representative confocal images of GATA1-stained HSCs isolated from control mice or 24hr after 450cGy X-ray irradiation (n=3). **(d)** Quantification of fraction of GATA1^+^ cells and GATA1 integrated density in HSCs isolated from control mice or 24hr after 450cGy x-ray irradiation(n (n=3, unpaired t-test for fraction of GATA1, and Mann-Whitney test for GATA1 IntDens). **(e)** Representative confocal images of HSCs isolated 24 hours after PBS or polyI:C treatment and stained for RPA32, γH2A.X, and DAPI. **(f)** Quantification of **(e)** for RPA32 and γH2A.X integrated density, and % RPA32^+^ cells. (n=3 independent sorts, each point an individual cell for RPA32 and γH2A.X integrated density, each point one individual experiment for % RPA32^+^ cells).

**ED2:**
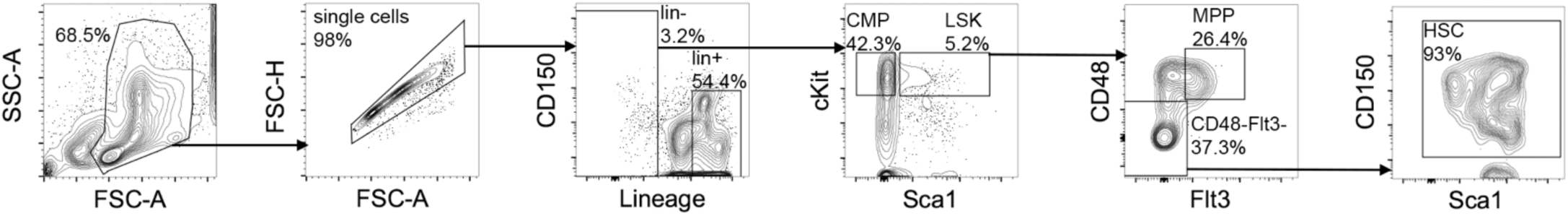
Sort gates for analysis of HSCs, MPPs, CMPs and lineage+ cells. Representative flow cytometry plots of gating strategy for sorting of mouse HSCs, MPPs, CMPs and lin^+^ cells.

**ED3:**
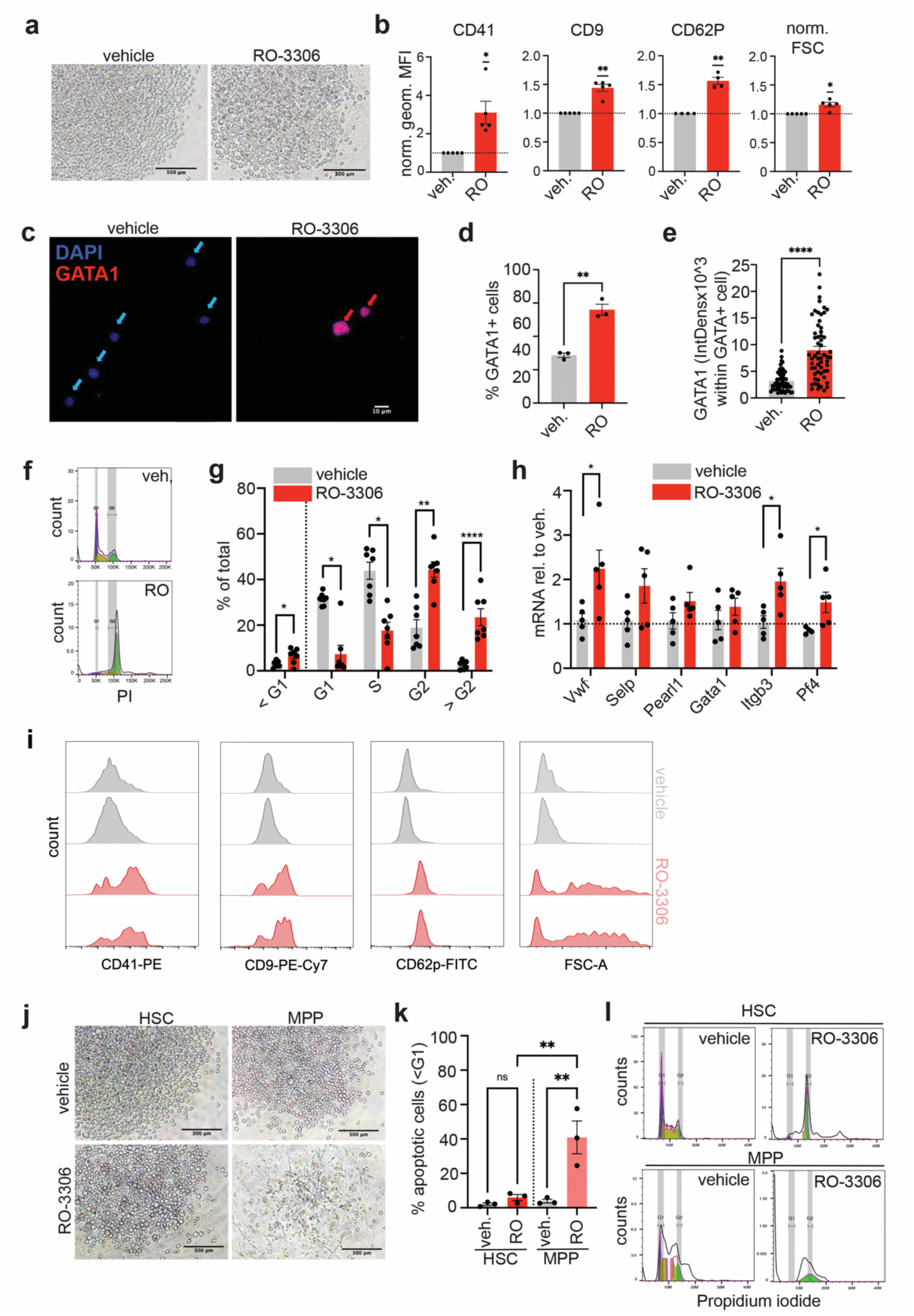
G2 arrest induces direct megakaryopoiesis from HSCs. a. Representative brightfield images of 48hr HSC cultures treated with 20mM RO-3306 (n=5). **b**. Flow cytometric analysis of MK surface markers in 20mM RO-3306– or vehicle-treated 48hr HSC cultures (n=5, each experiment averaged across technical replicates and normalized to vehicle, one-sample t-test). **c**. Representative confocal images of GATA1-stained HSCs after 48hrs of culture in the presence of 20mM RO-3306– or DMSO vehicle (n=3). **d**. Fraction GATA1^+^ cells in 48hr-cultured HSCs with RO-3306 or DMSO vehicle (n=3, unpaired t-test). **e**. Integrated density of GATA1 in 48hr HSC cultures with RO-3306 or DMSO vehicle (n=3, Mann-Whitney test). **f**. Flow cytometry histograms of cell cycle/ploidy analyses in 48hr DMSO vehicle or RO-3306-treated HSC cultures (representative of 4 experiments). **g**. Quantification of cell cycle/ploidy analyses in 48hr DMSO vehicle and RO-3306-treated HSC cultures (n=4, cell cycle stage determined with Watson’s pragmatic model and designated G1 and G2 ranges, paired t-tests). **h**. RT-qPCR for MK markers in 48hr DMSO vehicle– or RO-3306-treated HSC cultures (n=5, normalized to vehicle, unpaired t–test). **i**. Flow cytometry histograms of MK marker expression and FSC in 96hr DMSO vehicle and RO-3306-treated HSC cultures (representative of 4 experiments). **j**. Representative brightfield images of 48hr HSC and MPP cultures treated with DMSO vehicle or 20mM RO-3306. **k**. Fraction apoptotic cells as determined by PI cell cycle analysis in 48hr HSC and MPP cultures treated with DMSO vehicle or 20mM RO-3306 (n=3, patied t-tests). **l**. Flow cytometric histograms of cell cycle analysis in 48hr HSC and MPP cultures treated with DMSO vehicle or RO-3306 (representative of 3 experiments). *p ≤ 0.05, **p ≤ 0.01, ***p ≤ 0.001, and ****p ≤ 0.0001.

**ED4:**
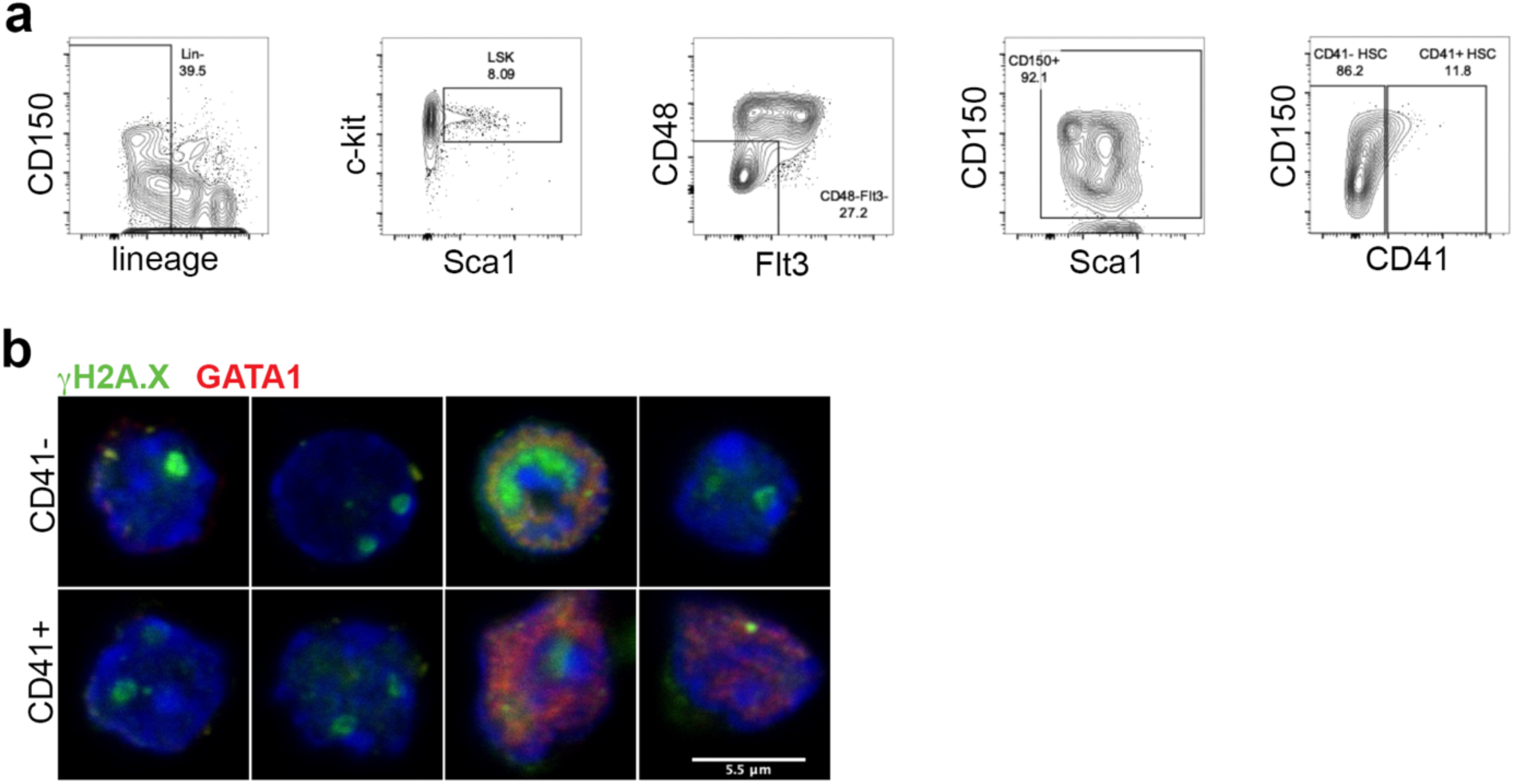
DNA damage in CD41^-^ and CD41^+^ HSCs a. Representative flow cytometry plots of gating strategy for sorting of mouse CD41^-^ and CD41^+^ HSCs. **b**. Representative confocal images CD41 and CD41 HSCs stained for γH2A.X and GATA1 (representative of 3 experiments).

**ED5:**
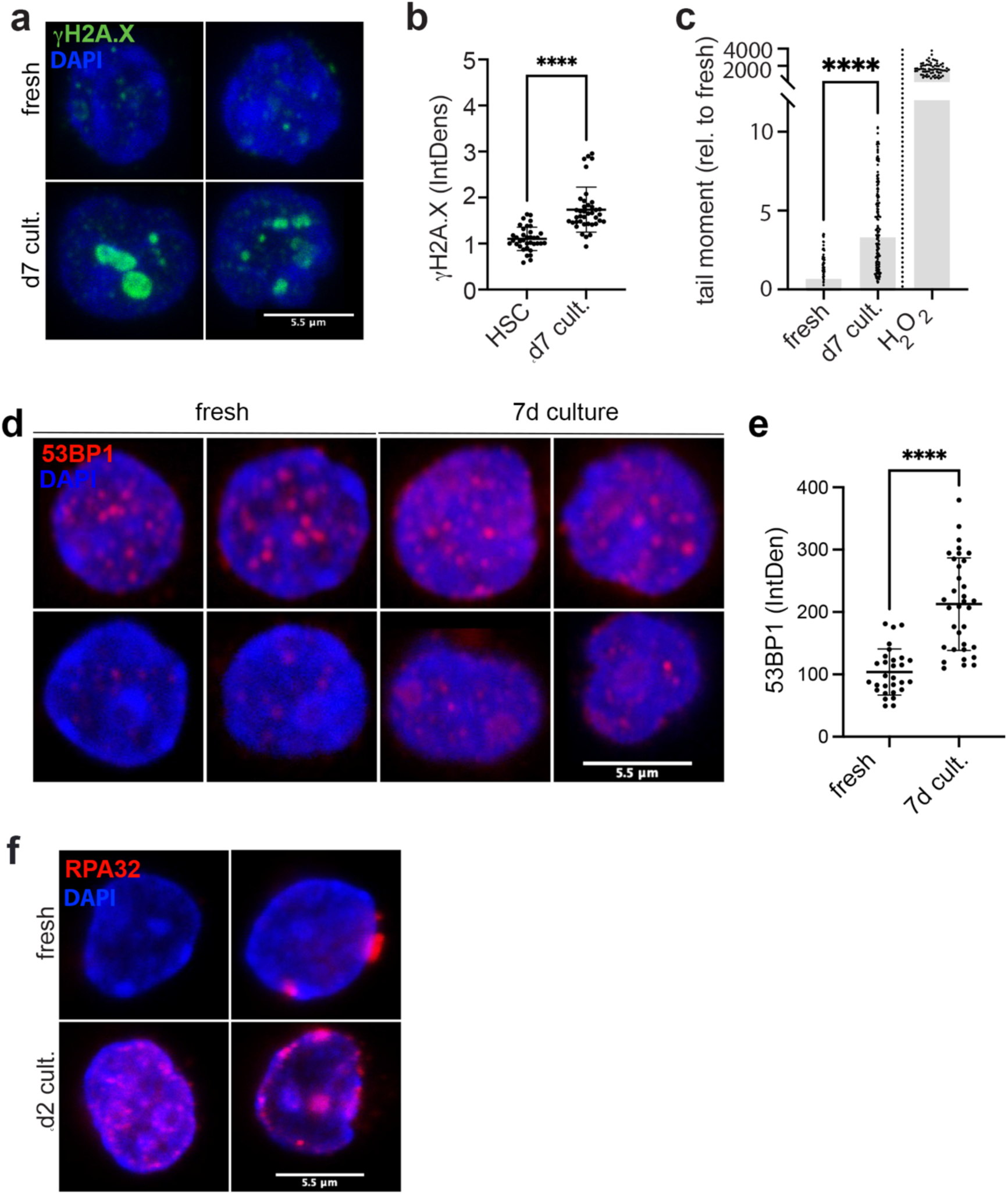
DNA damage in HSCs *in vitro*. a. Representative images of fresh HSCs or HSCs cultured for 7 days stained for γH2A.X (representative of 3 experiments). **b**. Quantification of fresh vs. d7 HSC cultures (n=3) for γH2A.X (Mann-Whitney test, n = 3). **c**. Comet assay of fresh and d7 cultured HSCs (H_2_O_2_-treated whole BM as positive control, n=3, Mann-Whitney test). **d**. Representative images of fresh HSCs or HSCs cultured for 7 days stained for 53BP1 (representative of 3 experiments). **e**. Quantification of 53BP1 deposition in fresh HSCs or HSCs cultured for 7 days (Mann-Whitney test, n = 3)**. (f)** Representative confocal images of fresh and 48hr-cultured HSCs stained for RPA32 (n=1).

**ED6:**
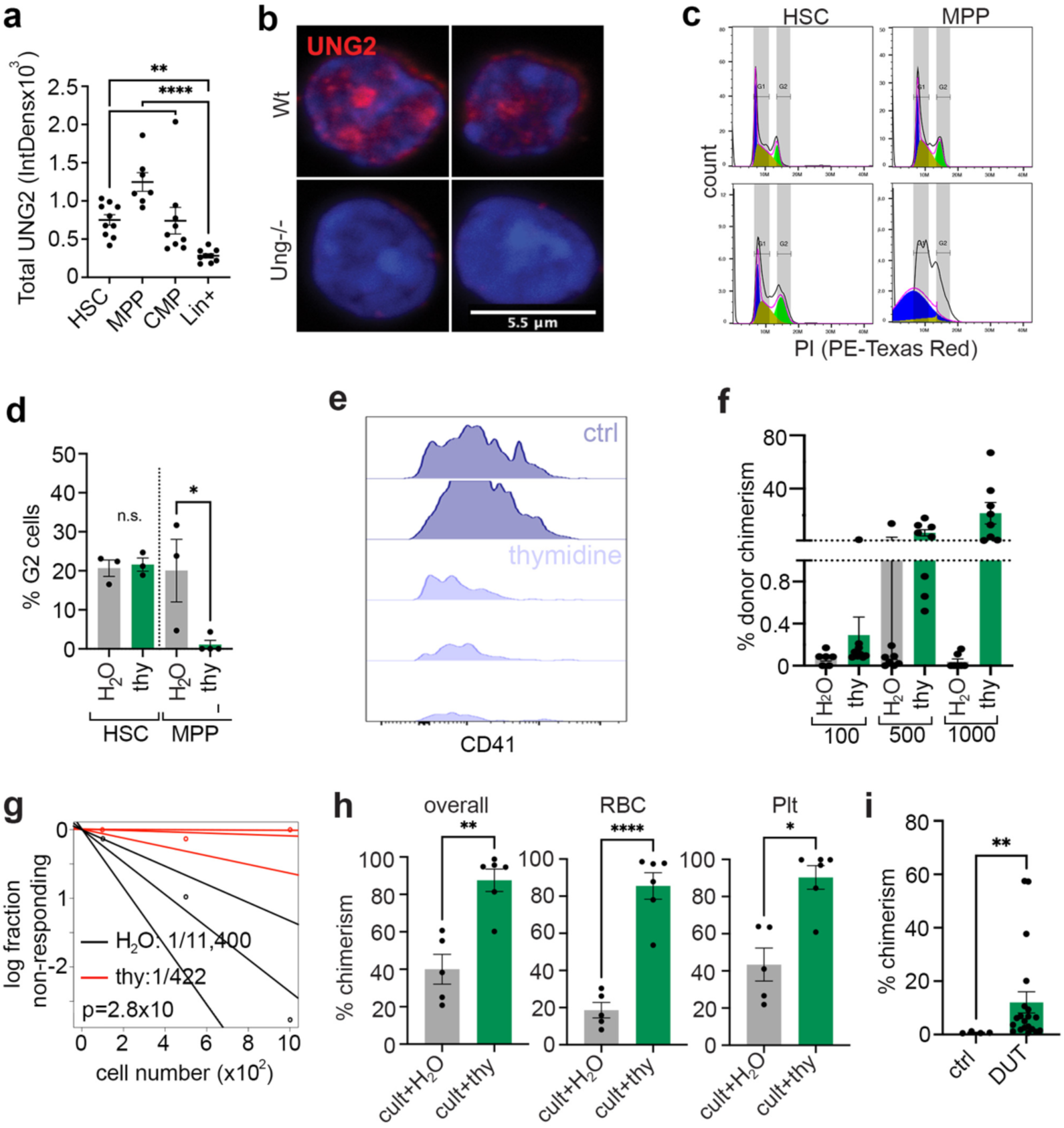
Uracil misincorporation and effect to thymidine. a. Quantification of nuclear UNG2 IntDens in fresh BM populations (n=1, one-way ANOVA). **b**. Representative confocal images of UNG2 staining in wt and *Ung^-/-^* HSCs, demonstrating specificity of the staining. **c**. Flow cytometry histograms of cell cycle/ploidy analyses in 48hr vehicle and thymidine-treated HSC and MPP cultures (representative of 3 experiments). **d**. Quantification of cell cycle/ploidy in 48hr vehicle or thymidine-treated HSC or MPP cultures (n=3, >1 technical replicate per experiment, multiple Mann-Whitney tests). **e**. Flow cytometry histograms of CD41 expression on phenotypic HSCs after 14 days of culture with vehicle or thymidine (representative of 6 experiments). **f**. Quantification of limiting dilution assay after transplantation of HSCs cultured with or without thymidine for 14d (positive chimerism > %1 donor derived). **g**. PB chimerism in individual recipients in limiting dilution assays after transplantation of HSCs cultured with vehicle or thymidine for 14d. **h**. KuO^+^ total, platelet, and RBC PB chimerism 16 weeks after competitive transplantation of 14d HSC cultures with vehicle or thymidine (n=5, unpaired t-test). *p ≤ 0.05, **p ≤ 0.01, ***p ≤ 0.001, and ****p ≤ 0.0001**. i**. 16-week post-competitive transplantation chimerism of HSCs transduced with *Dut* and cultured for 7 days (n= 8 recipients from two transplants, unpaired t-test).

